# Identification of reference genes for real-time PCR gene expression studies during seed development and under abiotic stresses in *Cyamopsis tetragonoloba* (L.) Taub

**DOI:** 10.1101/313437

**Authors:** Poonam Subhash Jaiswal, Navneet Kaur, Gursharn Singh Randhawa

## Abstract

Guar (*Cyamopsis tetragonoloba*) is an important industrial crop. The knowledge about genes of guar involved in various processes can help in developing improved varieties of this crop. qRT-PCR is a preferred technique for accurate quantification of gene expression data. This technique requires the use of appropriate reference genes from the crop to be studied. Such genes have not been yet identified in guar. The expression stabilities of the 10 candidate reference genes, viz., *CYP*, *ACT 11*, *EF-1α*, *TUA*, *TUB*, *ACT 7*, *UBQ 10*, *UBC 2*, *GAPDH* and *18S rRNA* were evaluated in various tissues of guar under normal and abiotic stress conditions. Four different algorithms, geNorm, NormFinder, BestKeeper and ∆Ct approach, were used to assess the expression stabilities and the results obtained were integrated into comprehensive stability rankings. The most stable reference genes were found to be *CYP* and *ACT 11*(tissues), *ACT 11*, *UBC 2* and *ACT 7* (seed development), *ACT 7* and *TUB* (drought stress), *TUA*, *UBC 2* and *CYP* (nitrogen stress), *TUA* and *UBC 2* (cold stress), *GAPDH* and *ACT 7* (heat stress) and *GAPDH* and *EF-1a* (salt stress). These results indicated the necessity of identifying a suitable reference gene for each experimental condition. Four selected reference genes were validated by normalizing the expression of *CtMT1* gene. To the best of our knowledge this is the first report on the identification of reference genes in guar. These findings are likely to provide a boost to the gene expression studies in this important crop.

## Introduction

Guar or clusterbean (*Cyamopsis tetragonoloba*, L. Taub.) is a self-pollinated and drought-tolerant legume crop. It is cultivated in the arid and semiarid regions of India, Pakistan, United States, Sudan, Brazil and Australia. The traditional use of guar is as a vegetable, cattle-feed and green manure crop. After the Second World War, guar has emerged as an industrial crop because of the industrial applications of galactomannan, commonly known as guar gum, which is present in the endosperm of its seeds. Guar gum, being a cost-effective natural thickner, binder and stabilizer, is an important raw material in a wide range of industries like petroleum, gas, food, paper, textile, cosmetic, mining and explosives (Dhugga et al. 2004). Guar gum has also shown a potential as a drug in the treatment of diseases like high cholesterol (Hosobuchi et al. 1999) diabetes (Saeed et al. 2012) and irritable bowel syndrome (Russo et al. 2015).

Despite the economic importance of guar, very few molecular genetic studies have been done on this crop. The major molecular genetic studies on guar are the cloning of mannan synthase gene (Dhugga et al. 2004), characterization of the mannan synthase promoter (Naoumkina et al. 2011) and guar leaf transcriptome profiling (Tanwar et al. 2017). Whole genome sequencing has not been yet done for guar. As a result of the limited information on molecular genetics of guar, the progress in the identification and characterization of the genes involved in important processes of this crop has been restricted. Gene expression analysis in guar can help in gaining insight into the functions and regulation of the genes involved in the growth and development, gum synthesis, and stress tolerance in this important crop plant. However, the lack of knowledge about stable reference genes has limited the gene expression studies in guar.

Gene expression analysis is carried out by various techniques like Northern blotting, semi-quantitative RT-PCR, cDNA microarrary, transcriptome profiling and quantitative real time PCR (qRT-PCR). The qRT-PCR technique is generally preferred due to its accuracy, specificity, sensitivity and less experimental time. The efficiency of qRT-PCR is decreased by the non-specific variations among cDNA samples. These variations are caused by the differences in RNA quality, reverse transcription efficiency and PCR efficiency. Hence, a normalization method is required for obtaining reliable and unbiased quantitative gene expression profiling of a target gene. Such a method involves the use of a reference gene(s). The reliability of the quantitative results of qRT-PCR depends on the selected reference gene(s). An ideal reference gene should have a stable expression in different tissues under different experimental conditions. A reference gene is generally a house-keeping gene (HKG) which is presumed to express consistently in various tissues under different environmental conditions. The house-keeping genes are involved in cellular maintenance to regulate basic and ubiquitous metabolic functions/pathways such as synthesis of the components of cytoskeleton, glycolytic pathway, synthesis of ribosome subunits, synthesis of elongation factors, protein folding and protein degradation. These genes are commonly used as reference genes assuming their stable expression in various tissues. However, recent research findings in several plant species, e.g., poplar hybrid (*P. trichocarpa* x *P. deltoidies*) (Brunner et al. 2004), *Arabidopsis thaliana* (Czechowski et al. 2005), *Solanum tuberosum* (Nicot et al. 2005), *Glycine max* (Jian et al. 2008), *Solanum lycopersicum* (Exposito-Rodriguez et al. 2008)*, Triticum aestivum* (Paolacci et al. 2009), *Oryza sativa* (Narsai et al. 2010), *Nicotiana benthamiana* (Liu et al. 2012), *Zeas mays* (Manoli et al. 2012), *Brassica juncea* (Chandna et al. 2012), *Arachis hypogea* (Reddy et al. 2013), *Vitis vinifera* (Borges et al. 2014), *Pennisetum glaucum* (Reddy et al. 2015), *Apium graveolens* (Li et al. 2016) have shown that the expression of the HKGs is not as stable as it is generally assumed. It is risky to randomly choose any HKG to serve as a reference gene for qRT-PCR. Therefore, the selection of an appropriate reference gene(s) is a crucial and challenging task in any gene expression study by qRT-PCR. As per our knowledge, no such study has been done in guar so far. In view of this, the present work was undertaken to identify the suitable reference genes for qRT-PCR gene expression studies in this industrially important crop.

## Materials and methods

### Plant material and treatments

The seeds of the guar variety HG-365 were obtained from Chaudhary Charan Singh Haryana Agricultural University (CCSHAU), Hisar, India. This is an early maturing, drought-tolerant and high yielding variety which has a high gum content. For surface sterilization, 100 seeds were treated with 0.1% (w/v) HgCl_2_ solution for 1-2 min, washed 4 times with sterilized distilled water and pre-soaked in 10 ml distilled water for 2 hours. The seeds were sown in pots containing sand and soil mixture (1:5) and the pots were placed in a screenhouse at the Indian Institute of Technology Roorkee, India. The soil moisture content was maintained close to the field capacity (10 ± 0.5%).

The experimental samples were classified into seven sets, namely, tissue set, seed development set, drought stress set, nitrogen stress set, cold stress set, heat stress set and salinity stress set, as per the requirement of the experiment.

### Tissue set

Young leaves, mature leaves, young roots, mature roots, nodules at 27 days after sowing (DAS), 38 DAS and 52 DAS, and developing seeds (35 days after flowering) were collected.

### Seed development set

Developing seeds at 15, 20, 25, 30, 35, and 40 days after flowering (DAF) were taken.

### Drought stress set

After 5 weeks, the water stress was given to the guar plants by withholding water until the plant reached permanent wilting point (soil moisture content 3.5±0.5%). The control plants were kept at the field capacity (soil moisture content 10±0.5%). The leaf and root samples were collected from drought treated and control plants.

### Nitrogen stress set

Guar seedlings were grown in full strength Hoagland medium. After 7 days, nitrogen stress was given to the guar seedlings by transferring them to nitrogen free Hoagland medium in hydroponics conditions. These plants were labelled as experimental plants. The guar seedlings in full strength Hoagland medium were labelled as control plants. The root and shoot tissue samples of control and experimental plants were taken at 2^nd^, 4^th^, 6^th^ and 8^th^ day post 7 days growth in Hoagland medium.

### Cold stress set

For cold stress treatment, the pots with the 7-day-old seedlings were kept in a growth chamber at 4 °C for 0, 2, 6, 12 and 24 h. The leaf and root samples were collected from each treatment.

### Heat stress set

For heat stress treatment, the pots with the 7-day-old seedlings were kept in a growth chamber at 42 °C for 0, 2, 6, 12 and 24 h. The leaf and root samples were collected from each treatment.

### Salinity stress set

The salt stress was given to 7-day-old seedlings for 0, 6, 12, 24, 48 and 72 h by completely saturating the potted plants with 120 mM NaCl solution. The leaf and root samples were collected from each treatment.

The tissue samples from each experimental set were collected in triplicate, quickly frozen in liquid nitrogen and stored in a deep-freezer at −80°C till RNA isolation.

## Selection of candidate reference genes and primer designing

The HKGs, namely, cyclophilin *(CYP)*, actin 11 *(ACT 11)*, actin 7 *(ACT 7)*, elongation factor -1 alpha *(EF-1a)*, alpha-tubulin *(TUA)*, beta-tubulin *(TUB)*, polyubiquitin 10 *(UBQ 10)*, ubiquitin conjugating enzyme E2 2 *(UBC 2)*, glyceraldehyde-3-phosphate dehydrogenase (*GAPDH*) and 18S ribosomal RNA *(18S rRNA)* were selected to analyse the expression stability of these genes in different experimental conditions by using qRT-PCR. The sequences for first nine selected genes of different legumes were retrieved from NCBI database. Multiple sequence alignment of sequences of each gene from different legumes was performed by Clustal Omega to identify conserved regions of a particular gene. The primer pair for each selected gene was designed manually from the conserved regions and used in PCR to amplify the desired gene of guar. The sequencing of the PCR products was outsourced to Chromas Biotech. Pvt. Ltd., Bengaluru (India). The sequence data obtained from the company were confirmed by the nucleotide BLASTN tool and submitted to NCBI. The NCBI accession numbers of the submitted partial cDNA sequences are MF370613 *(CYP)*, MF370605 *(ACT 11)*, MF370611 *(EF-1α)*, MF370607 *(TUA)*, MF370608 *(TUB)*, MF370606 *(ACT 7)*, MF370610 *(UBQ 10)*, MF370612 *(UBC 2)* and MF370609 *(GAPDH)* (unpublished).

The primer pairs for 9 reference genes, viz., *CYP*, *ACT 11*, *ACT 7*, *EF-1a*, *TUA*, *TUB*, *UBQ 10*, *UBC 2* and *GAPDH*, were designed from the submitted sequence data of the genes of guar for qRT-PCR studies. The primer pair for *18S rRNA* reference gene was designed on the basis of the sequence of this gene obtained from NCBI (JX444503). The designed primers were checked by OligoAnalyzer tool. The various parameters of each primer pair were as follows: melting temperature 61-67°C, primer length 20-26 bp, GC content between 45-60%, and the PCR product length 93-205 bp. The target gene sequences were amplified by PCR using the designed primers and the amplicons obtained were confirmed by sequencing. The characteristics of primer sequences used in this study are given in Table 1.

## Total RNA isolation and synthesis of cDNA

Total RNA from a guar plant sample was isolated with HIMEDIA^®^ RNA-XPress reagent according to the protocol given by the manufacturer. The isolated RNA was given an RNase-free DNase I (Thermo scientific) treatment following manufacturer’s instructions to remove DNA contamination in the sample. The concentration and purity of the isolated total RNA were determined using a DeNovix DS-11 spectrophotometer. The RNA samples having the absorption ratios of A260/A280 = 1.9 to 2.08 and A260/A230 = 2.04 to 2.18 were used for synthesis of cDNA. The quality of total RNA was verified by performing 1.5% agarose gel electrophoresis. An aliquot of 1μg of a total RNA sample was used for first-strand cDNA synthesis using verso cDNA synthesis kit (Thermo scientific) for the 20 μl reaction mixture following the manufacturer’s protocol with slight modification in the reverse transcription cycling program (1 cycle of 60 min at 42 °C and 2 min at 95°C).

## Quantitative Real-Time PCR

qRT-PCR was performed according to the Minimum Information for Publication of Quantitative Real-Time PCR Experiments (MIQE) guidelines (Bustin et al. 2009) in a 96-well plate using StepOnePlus Real-Time PCR (Applied Biosystems) following manufacturer’s instructions. Power SYBR Green dye (Applied Biosystems) was used to measure the cycle threshold (Ct) or quantification cycle (Cq) value. The quantitative real-time PCR program was set to 2 min at 50 °C, 10 min at 95 °C, and 15 s at 95 °C and 1 min at 60 °C (40 cycles) as described in Sinha et al. (2015). The melt curve was added in the above program to check the amplicon specificity. The PCR efficiency (E) for each candidate reference gene was calculated from the slope of the standard curve based on 10-fold serial dilution of pooled cDNA over five dilution points (Sinha et al. 2015) and correlation coefficient (R^2^) for each gene was also determined. For expression profiling of a candidate reference gene, the reaction was performed in 3 biological and 2 technical replicates along with a no template control (NTC).

## Analysis of the expression stability of reference genes

The expression stability of a selected reference gene was evaluated by the following four methods.

### Stability analysis by qbase+ tool

The qbase+ tool is based on geNorm (Vandesompele et al. 2002) and qBase (Hellemans et al. 2007) technologies. The average expression stability value (*M*) of a reference gene was determined by geNorm analysis using qbase+ tool according to the instructions given in the qbase+ manual (www.qbaseplus.com). According to the geNorm analysis, the most stable reference gene is the one having the lowest *M*-value. The geNorm analysis also calculates pair-wise variation (Vn/n+1) and determines the number of reference genes required for normalizing the gene expression data. Vn/n+1 ratio below 0.15 suggests that the use of an additional reference gene would not significantly improve normalization (Vandesompele et al. 2002).

### Stability analysis by NormFinder tool

The NormFinder is an algorithm for identifying suitable reference genes for normalization on the basis of stability value (SV) (Andersen et al. 2004). The SV for each candidate reference gene was determined following the software guidelines (https://moma.dk/normfinder-software). As per the NormFinder analysis, the candidate reference gene having the lowest SV is the most stable reference gene.

### Stability analysis by BestKeeper tool

The stability analysis of a candidate reference gene was performed according to the instructions of the BestKeeper tool (www.gene-quantification.com/bestkeeper.html). The BestKeeper tool ranks the stabilities of candidate reference genes on the basis of standard deviation (SD) and coefficient of variation (CV) values of the studied genes (Pfaffl et al. 2004). The most stable reference gene is the one having lowest SD value.

### ∆Ct approach

The expression stability of a candidate reference gene was determined by calculating the ΔCt value and standard deviation (StdDev) as described in Silver et al. (2006) and Jian et al. (2008). As per the analysis, the candidate reference gene having the lowest mean standard deviation is the most stable reference gene.

### Comprehensive ranking

The comprehensive ranking of expression stability of a candidate reference gene was determined by calculating the geometric mean of four types of rankings obtained from geNorm, NormFinder, BestKeeper and ∆Ct approach according to the RefFinder approach (Stajner et al. 2013) as described in Xiao et al. (2015).

### Validation of selected reference genes

The selected reference genes in different tissues and under drought stress condition were validated by using *Cyamopsis tetragonoloba* metallothionein-like type 1 (*CtMT1*) gene (GenBank accession no. **KU903285**) as a target gene. The relative expression of *CtMT1* was analysed by using most stable genes (*CYP* and *ACT 11*) and least stable gene (*TUB*) in tissue samples. For drought stress, *ACT 7* & *TUB* (most stable genes) and *18S rRNA* (least stable gene) were used for normalization. qRT-PCR reaction was performed same as that of described earlier in material and method section. The relative changes in *CtMT1* expression levels were calculated by using the 2^−ΔCt^ (Schmittgen and Zakrajsek 2000) method in different tissues and 2^−ΔΔCt^ method (Livak and Schmittgen 2001) in drought stress conditions as described in Feng et al. (2018).

## Results

### Amplification specificities and PCR efficiencies of reference genes

The lengths of the PCR amplification products of 10 candidate reference gene fragments ranged from 93 to 205 bp (Table1 and Supplementary Figure S1). Each amplicon gave a single band on an agarose gel (Supplementary Figure S1) and its length matched with amplicon size of the selected fragment of corresponding gene as given in Table 1. The sequence of this amplified product was also identical with the corresponding guar gene sequence (data not shown). This confirmed that the primer pair amplified a unique cDNA fragment corresponding to the selected candidate reference gene. The PCR efficiencies (E) and the correlation coefficients (R^2^) for the 10 candidate reference genes ranged from 95.91 to 103.53 and 0.998 to 1.000, respectively (Table 1). These results were in the acceptable range of both PCR efficiency and correlation coefficient as per the MIQE guidelines (Bustin et al. 2009) indicating that the qRT-PCR amplifications were quite good.

## Expression profiles of reference genes

The expression profiles of 10 candidate reference genes, viz., *CYP*, *ACT 11*, *EF-1α*, *TUA*, *TUB*, *ACT 7*, *UBQ 10*, *UBC 2*, *GAPDH* and *18S rRNA* in different samples of guar have been presented in Figure 1. The expression level of each candidate reference gene in a sample has been represented as a quantification cycle value (Cq). The Cq values of the 10 candidate reference genes in various samples were in a range of 7.64 to 33.19 (Figure 1 and Supplementary Table S1). The mean Cq values of reference genes varied from 10.86 for *18S rRNA* to 28.85 for *UBQ10* (Supplementary Table S1). The range of Cq values for a gene represents the variation in the expression of this gene in 66 samples. The Cq values showed a low variation for the genes *UBC 2*, *ACT 7*, *ACT 11*, *CYP* and *TUA* which indicated that there was a low variation in the expression of these genes in different guar plant samples. High variation in the Cq values was observed for the genes *TUB*, *18S rRNA*, *UBQ10 GAPDH*, and *EF1-α* (Figure 1), showing high variation in the expression of these genes in different guar samples.

**Figure 1.**
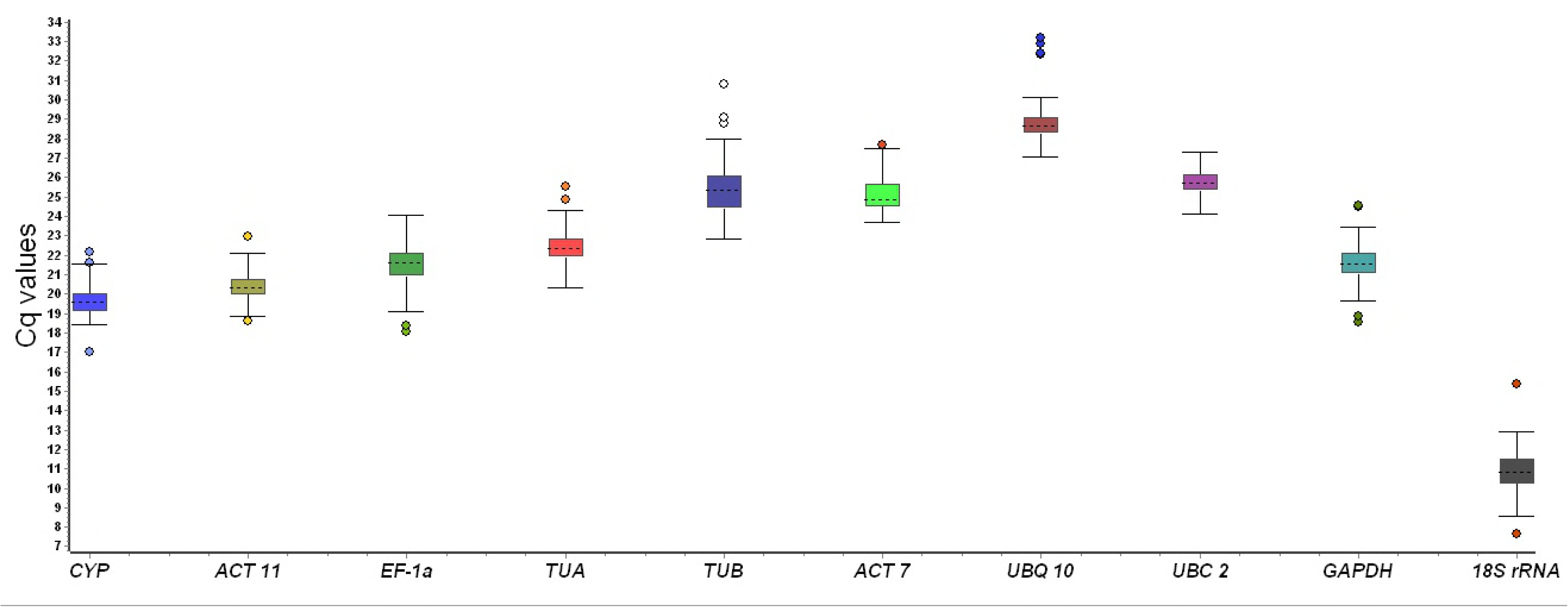
Expression levels of the candidate reference genes in all samples. Median, interquartile range, non-outlier range and outlier for each gene have been depicted in the box plots.

## Stability analysis of candidate reference genes expression

The results of expression stabilities of the selected candidate reference genes in different tissues at various growth stages and under different abiotic stresses, as determined by four algorithms have been given below.

## geNorm analysis

The average expression stability rankings of the 10 candidate reference genes in seven experimental sets by geNorm analysis have been given in Figure 2 and Supplementary Table S2. According to the geNorm analysis by qbase+ software, the gene(s) having the lowest *M-*value is most stable and the gene(s) having the highest *M-* value is least stable. The *M-*value of each of the 10 candidate reference genes was less than the 1.5 cut-off value (Vandesompele et al. 2002). The lowest *M* values in the tissue set (*M* = 0.094) and nitrogen stress set (*M* =0.395) were observed for the *CYP* gene and hence this was the most stable reference gene in these sets. The *UBC 2* gene having the lowest *M-*value was the most stable reference gene in the seed development set (*M* value of 0.462) and in cold stress set (*M-*value of 0.155). The *ACT 7* gene having lowest *M-*value of 0.29 in drought stress set and *M-*value of 0.113 in heat stress set, was the most stably expressed reference gene. The lowest *M* value in the salt stress set (*M* = 0.378) was observed for the *ACT 7* and *EF-1α* genes and hence these were the most stable reference genes in this set. The *TUB* gene was found to be least stable in tissue set. In seed development, nitrogen and cold stress sets, the least stable genes were *TUA*, *UBQ 10* and *ACT 7*, respectively. The *18S rRNA* was the least stable gene in drought, heat and salt stress sets.

**Figure 2.**
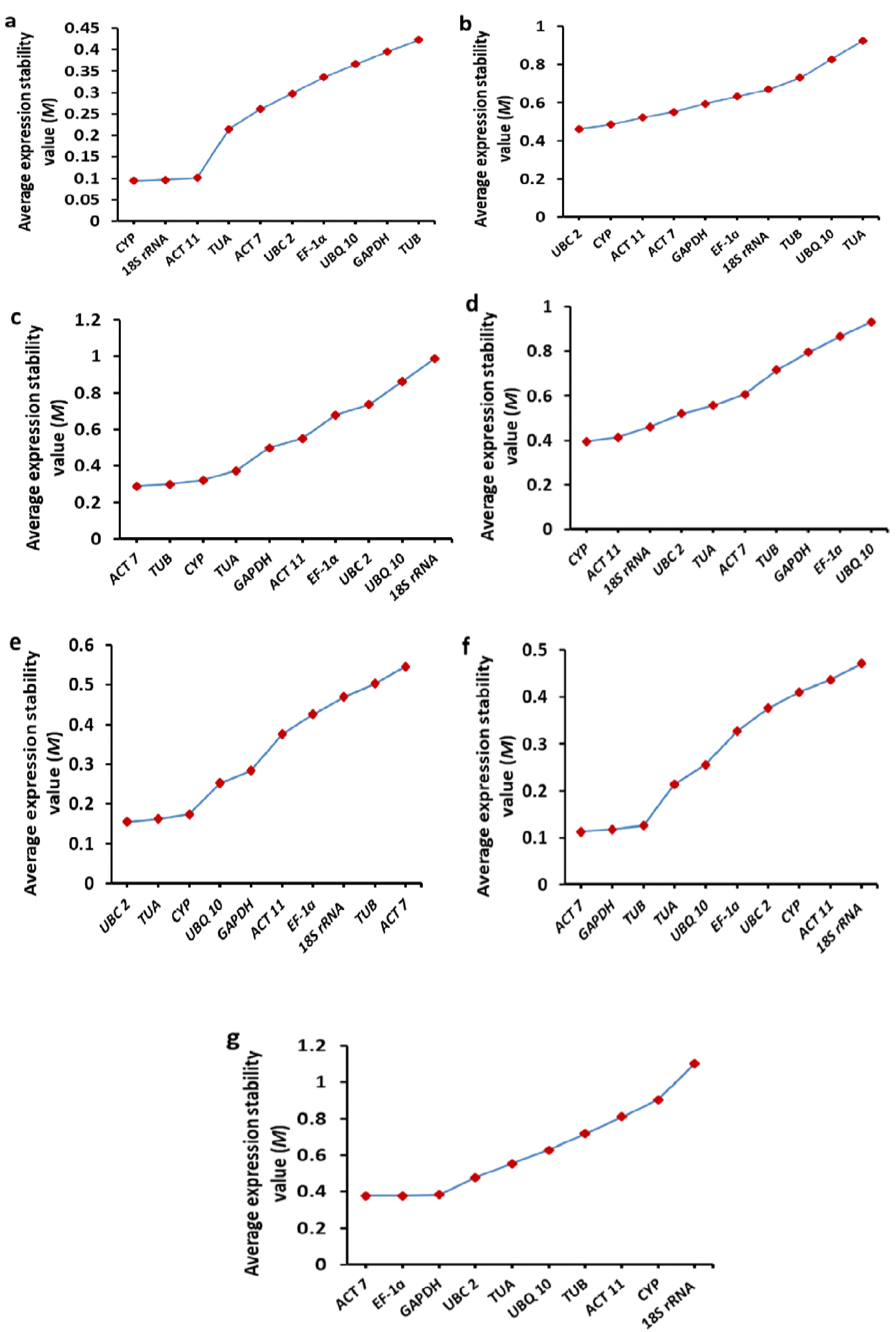
Average expression stability graphs of candidate reference genes by geNorm analysis. Average expression stability values of candidate reference genes in tissue set (a), seed development set (b) drought stress set (c), nitrogen stress set (d), cold stress set (e), heat stress set (f) and salinity stress set (g). A lower average expression stability (*M*) value indicates more stability in the expression of the candidate gene.

The results of pairwise variation (V) in seven experimental sets by geNorm analysis have been given in Figure 3. According to geNorm principle, the Vn/n+1 value less than 0.15 suggests that one more reference gene is not required (Vandesompele et al. 2002). The V 2/3 values for tissue, drought stress, cold stress, heat stress and salinity stress sets were 0.035, 0.117, 0.063, 0.046 and 0.115, respectively, indicating that the two most stable reference genes are enough for reliable normalization in each of these experimental sets and a third reference gene is not required. For seed development and nitrogen stress sets, V3/4 values were 0.124 and 0.127, respectively. Hence, three most stable reference genes are sufficient in these sets.

**Figure 3.**
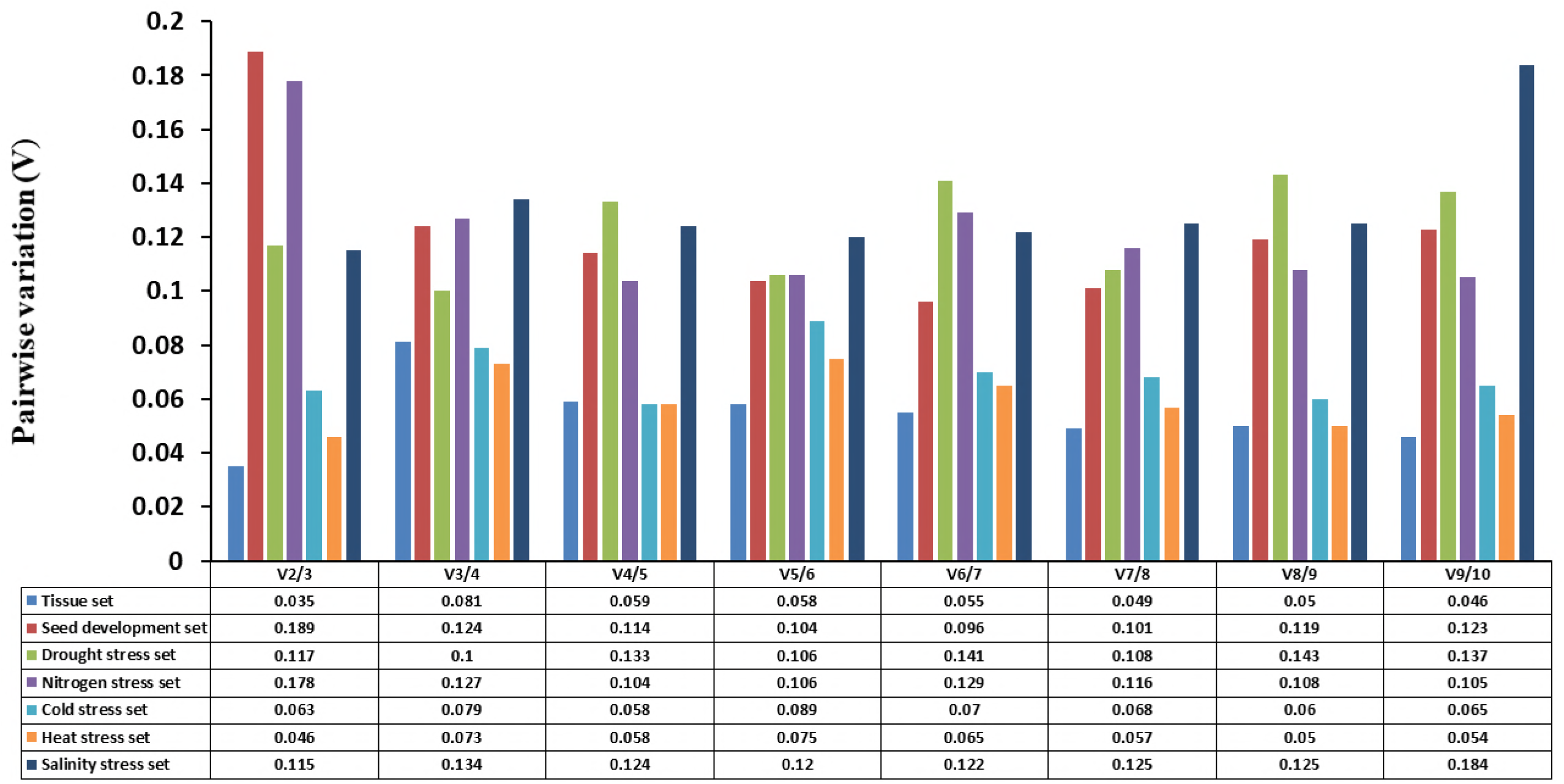
Pairwise variation (V) values in seven experimental sets using geNorm analysis. The Vn/n+1 value <0.15 indicates that one more reference gene is not required for normalization.

## NormFinder analysis

The stability values (SVs) obtained from the NormFinder analysis of the expression of 10 candidate reference genes in seven experimental sets are presented in Figure 4 and Supplementary Table S3. According to NormFinder software rule, the gene having the lowest SV is the most stable one for qRT-PCR. In the tissue set, the lowest SV (0.005) was observed for the *ACT 7* and *ACT 11* genes and hence these genes were most stable in this set. The *ACT 11* gene having the lowest SV (0.001) was the most stably expressed reference gene in seed development set. In the drought stress set, *TUB* and *ACT 7* genes had the lowest SV (0.001) and hence both these genes were the most stably expressed gene in drought stress condition. In the nitrogen stress set, the lowest SV (0.013) was observed for the *TUA* gene and hence this gene was most stable in this set whereas *TUA* and *CYP* genes were the most stably expressed reference genes in cold stress set. *UBC 2* and *GAPDH* genes were the most stable reference genes in heat stress set and salt stress set, respectively. The *18S rRNA* was the least stable gene in all the experimental sets (Figure 4).

**Figure 4.**
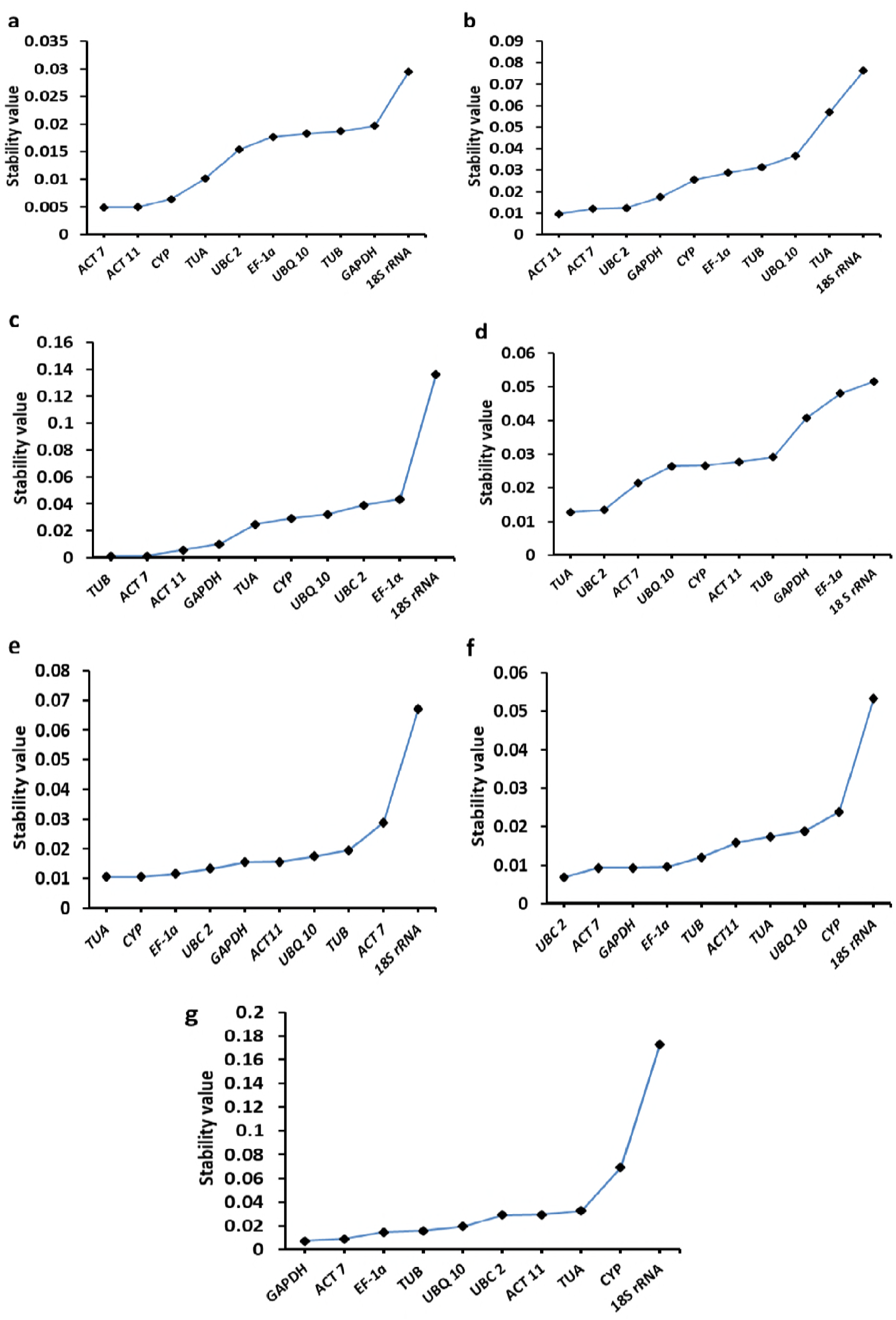
Expression stability graphs of candidate reference genes by NormFinder analysis. A lower stability value (SV) indicates more stable expression in tissue set (a), seed development set (b) drought stress set (c), nitrogen stress set (d), cold stress set (e), heat stress set (f) and salinity stress set (g).

## BestKeeper analysis

The standard deviation (SD) values obtained from the BestKeeper analysis of the expression of 10 candidate reference genes in seven experimental sets are presented in Figure 5 and Supplementary Table S4. According to the BestKeeper software rule, the candidate reference gene having the lowest SD value is the most stable reference gene. The presented data showed that the *UBC 2* gene expressed most stably in tissue set and drought stress set whereas the *CYP* gene was the most stable reference gene in seed development set. The *ACT 11* gene had the most stable expression in nitrogen stress set and *TUA* gene was the most stable reference gene in cold, heat and salt stress sets. (Figure 5).

**Figure 5.**
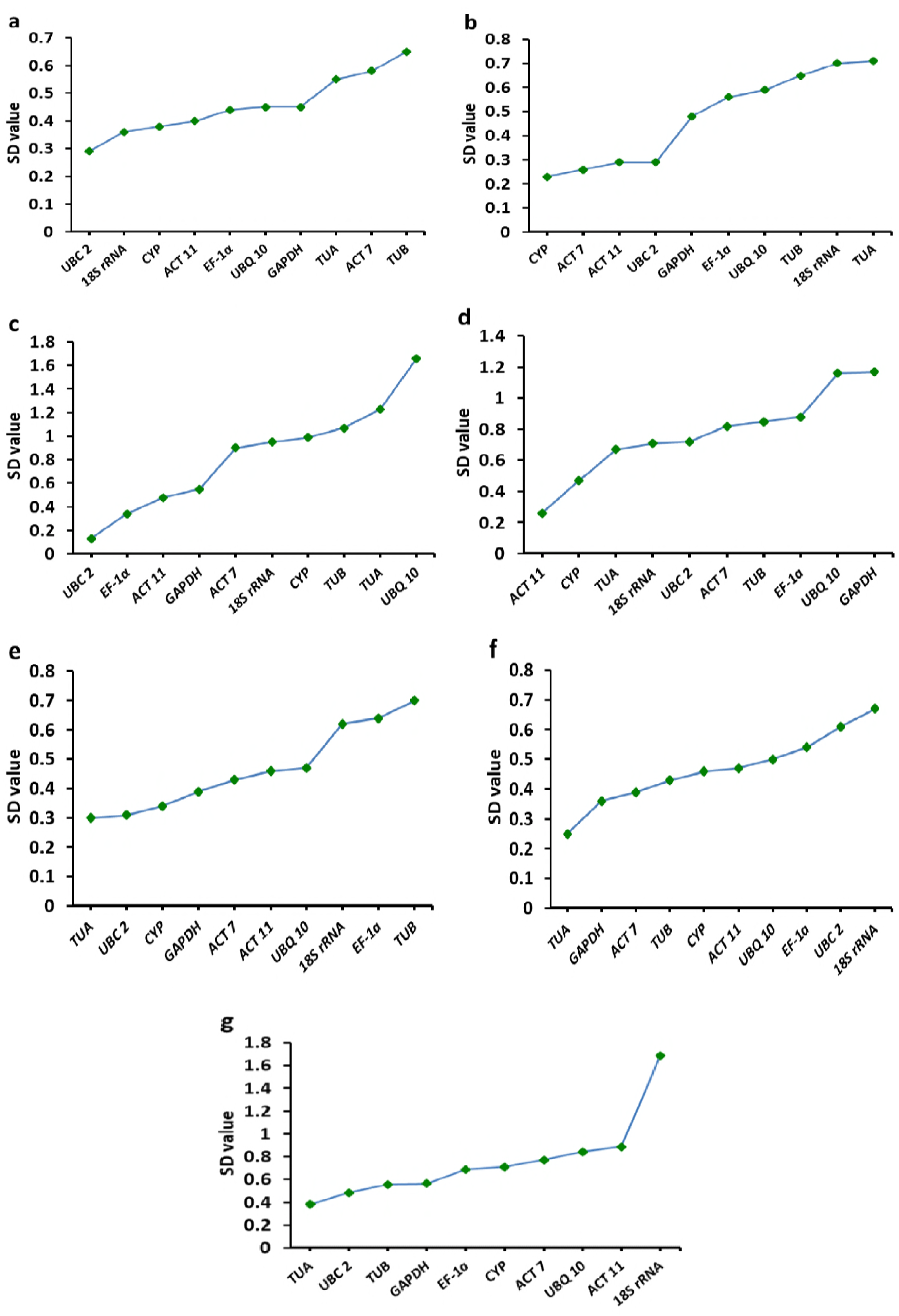
Expression stability graphs of candidate reference genes by BestKeeper analysis. A lower SD value shows higher stability of reference gene in tissue set (a), seed development set (b) drought stress set (c), nitrogen stress set (d), cold stress set (e), heat stress set (f) and salinity stress set (g). Abbreviation: SD - standard deviation.

## ΔCt analysis

The comparisons of the all possible combinations of the 10 reference genes in the seven experimental sets by ΔCt approach have been given in Supplementary Table S5 to S11. According to the ΔCt approach, a candidate reference gene having the lowest mean StdDev value is the most stable reference gene (Jian et al. 2008). Hence, the *18S rRNA* gene having the lowest mean StdDev value 0.32 in tissue set was the most stably expressed reference gene and the *TUB* having the highest mean StdDev value 0.53 was the least stable reference gene in this set (Table 2 and Supplementary Table S5). In seed development set, *ACT 11* was the most stable gene (mean StdDev, 0.72) and *UBQ 10* was the least stable gene (mean StdDev, 1.42) (Table 2 and Supplementary Table S6). The *ACT 7* (mean StdDev, 0.73) and *18S rRNA* (mean StdDev, 1.50) genes were the most stable and least stable genes, respectively in drought stress set (Table 2 and Supplementary Table S7). The *TUA* (mean StdDev, 0.74) gene was the most stable gene in nitrogen set (Table 2 and Supplementary Table S8). The *GAPDH* (mean StdDev, 0.38) and *TUA* (mean StdDev, 0.38) genes were the most stable genes in cold stress set (Table 2 and Supplementary Table S9). The most stable gene in heat stress set was *GAPDH* (mean StdDev, 0.34) (Table 2 and Supplementary Table S10). *GAPDH* and *EF-1α* genes had the lowest mean StdDev (0.83) and were hence the most stably expressed genes in salt stress set. In this set *18S rRNA* gene having the highest mean StdDev (1.90) was the least stable reference gene (Table 2 and Supplementary Table S11).

## Comprehensive ranking of expression stability

As described in the Materials and methods, the comprehensive ranking of the expression stabilities of 10 candidate reference genes was done by the RefFinder approach (Stajner et al. 2013). The Table 3 shows the comprehensive ranking of gene expression stabilities for 10 reference genes. The *CYP* gene ranked first and *ACT 11* gene ranked second in the tissue set, indicating that these two genes were the most stably expressed reference genes in this set. In seed development set, *ACT 11*, *UBC 2* and *ACT 7* genes being the first, second and third ranked genes, were the most stable reference genes. The *ACT 7* gene ranked first and the *TUB* gene ranked second in the drought stress set indicating that these genes had the most stable expression in this set. In nitrogen stress set, the *TUA* gene ranked first and, the *UBC 2* and *CYP* ranked second. Hence, these three genes were the most stably expressed reference genes under nitrogen stress condition. The *TUA* and *UBC 2* genes ranked first and second, respectively, in stability of expression in cold stress set. In heat stress set, *GAPDH* and *ACT 7* genes were on the first and second rank. *GAPDH* and *EF-1α*, the first and second ranked genes in salt stress set, were the most stable reference genes under this condition. The least stable reference gene in tissue and cold stress sets was *TUB* gene whereas *TUA* and *EF-1α* genes were the least stable genes in seed development and nitrogen stress sets, respectively. The *18S rRNA* gene was the least stable reference gene in drought, heat and salt stress sets.

## Validation of selected reference genes in different tissues

The relative expression levels of *CtMT1* in different tissues of guar using most stable and least stable genes for normalization have been presented in Figure 6a. When most stable reference gene, *CYP* or *ACT 11*, was used for normalization, the *CtMT1* gene was found to express at very low level in young leaves (YL) and mature leaves (ML) whereas its higher expression pattern was observed in young root (YR) and mature root (MR). However, when the least stable reference gene, *TUB* was used, the relative expression level of *CtMT1* gene was found to be high in YL and ML, and abnormally high in YR and MR.

**Figure 6.**
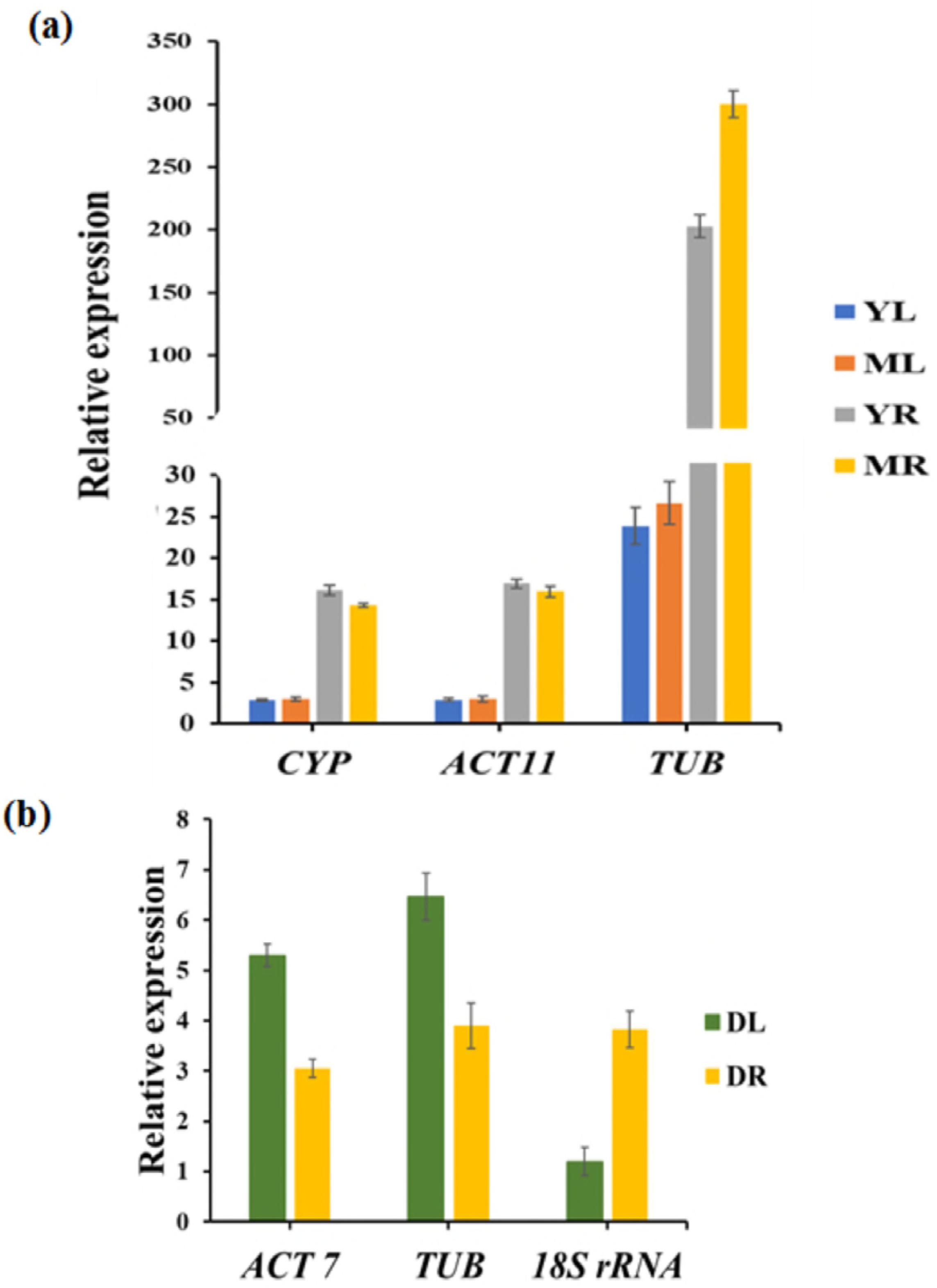
Relative expression level of *CtMT1* in (a) different tissues and (b) drought stress condition of guar. Most stable gene (*CYP* or *ACT 11* for different tissues and *ACT 7* or *TUB* under drought stress) and least stable gene (*TUB* for different tissues and *18S rRNA* under drought stress) were used as reference genes for normalization. The error bar indicates standard deviation (± SD) of three biological replicates. Abbreviation: Young leaves- YL, mature leaves- ML, young roots- YR, mature roots- MR, leaves under drought stress- DL, roots under drought stress- DR.

## Validation of selected reference genes during drought stress condition

The relative expression levels of *CtMT1* gene in leaves and roots of guar under drought stress condition using most stable and least stable genes for normalization have been presented in Figure 6b. In leaves, when most stable reference genes *ACT 7* and *TUB* were used, *CtMT1* showed upregulation up to 5.29 fold and 6.47 fold, respectively (Figure 6b), however, 1.19 fold change was observed when the least stable reference gene (*18S rRNA*) was used for normalization indicating that the expression pattern of *CtMT1* was found to be nearly unchanged in leaves under drought stress when least stable reference gene was used. Increase in the fold change up to 3.0 fold and 3.89 fold was observed for *CtMT1* in roots under drought condition when most stable reference genes *ACT 7* and *TUB*, respectively, were used, whereas the expression pattern of *CtMT1* was 3.83 fold when the least stable reference gene (*18S rRNA*) was used for normalization (Figure 6b). The above results indicated upregulation in the expression of *CtMT1* gene in both leaves and roots under drought stress condition when the most stable reference gene, *ACT 7* or *TUB* was used for normalization, however the level of upregulation was higher in leaves as compared to that in roots.

## Discussion

Guar is an important industrial crop and as a result of its increased demand there is a need to develop its improved varieties having more stress tolerance and producing increased amounts of high quality gum. The knowledge about the genes involved in development/growth, stress tolerance and seed/gum production will be very helpful to develop improved varieties of guar. Gene expression analysis is an appropriate approach to identify the genes and understand their functions and regulation in various important processes of guar crop. The qRT-PCR technique, which is often used to study gene expression, requires the use of a reference gene(s) for normalization. The present study is the first report on the selection of suitable reference genes of guar for gene expression studies in different tissues at various growth stages, seed development and different abiotic stress conditions.

Ten house-keeping genes, viz., *CYP*, *ACT 11*, *EF-1α*, *TUA*, *TUB*, *ACT 7*, *UBQ 10*, *UBC 2*, *GAPDH* and *18S rRNA* were selected for gene expression studies in various tissues of guar under different experimental conditions. In order to select the most suitable reference gene(s) in an experimental set, four different methods, viz., geNorm analysis, NormFinder analysis, BestKeeper analysis and ∆Ct approach, were used. Our results showed that the rankings obtained from these four methods were not generally identical. Hence, for this study, a most stable reference gene was selected on the basis of comprehensive ranking. Similar approach has also been reported in *Salicornia europaea* by Xiao et al. (2015) and in chickpea by Reddy et al. (2016).

Our study indicated that *CYP* and *ACT 11* genes were found to be most suitable reference genes for accurate normalization of the gene expression data in various tissues of guar under normal conditions. In the earlier studies, the *CYP* gene was also found to be a suitable reference gene for qRT-PCR normalization in different tissues of soybean (Jian et al. 2008) and peanut (Reddy et al. 2013). However, this gene was not a suitable reference gene in poplar (Brunner et al. 2004), cucumber (Wan et al. 2010) and chickpea (Reddy et al. 2016). Similarly, *ACT 11* gene was also not found to be a suitable reference gene for qRT-PCR studies in different tissues of poplar (Brunner et al. 2004). For the gene expression studies in developing guar seed at various stages, the *ACT 11*, *UBC 2* and *ACT 7* genes were found to be the most suitable reference genes. In a previous report, *ACT 7* gene was the most suitable reference gene for developmental stage gene expression study in soybean (Jian et al. 2008). However, this gene was not the suitable reference gene under various developmental stages in rice (Pabuayon et al. 2016) and *Lilium regale* (Liu et al. 2016).

Guar is normally cultivated in marginal soils where drought, salt, temperature and nutrient stresses are very common. qRT-PCR gene expression studies under these abiotic stress conditions in guar can help in undermining the molecular mechanisms of tolerance to these stresses. Our investigations revealed that *ACT 7* and *TUB* genes were most suitable as a reference for accurate normalization of the gene expression data under drought stress in guar. The *TUB* gene was also earlier found to be the most suitable reference gene under drought stress study in *Platycladus orientalis* (Chang et al. 2012), *Musa* (Podevin et al. 2012), maize (Lin et al. 2014), and pigeonpea (Sinha et al. 2015) but this gene was not a suitable reference gene in chickpea (Garg et al. 2010) and *Salicornia europaea* (Xiao et al. 2015). Similarly, *ACT 7* gene was not the proper reference gene under drought stress condition in *Brachypodium distachyon* (Hong et al. 2008) and *Platycladus orientalis* (Chang et al. 2012). Our findings would prove to be an anchor in the research work aiming to identify drought tolerant guar genes which may be used to make drought tolerant varieties not only of guar but of other crops too.

Guar, being a legume crop, is able to fulfill its nitrogen requirement through biological nitrogen fixation. However, this crop usually encounters nitrogen deficiency during the initial stages of crop growth when nitrogen fixing nodules have not been fully developed. Thus, it is important to study the mechanism of nitrogen uptake in guar seedlings. Gene expression studies under nitrogen stress have not been carried out yet in guar, due to unavailability of a suitable reference gene(s). Our study has revealed that *TUA*, *UBC 2* and *CYP* genes were the most suitable reference genes in guar under nitrogen stress condition. *TUA* gene was also found to express stably in *Salicornia europaea* under nitrogen stress treatment (Xiao et al. 2015). Our findings are likely to facilitate the research initiatives to determine the genes expressing under nitrogen stress in guar.

In the earlier studies, *GAPDH* and *ACT 7* genes were found to be suitable reference genes for qRT-PCR normalization under heat stress condition in *Brachypodium distachyon* (Hong et al. 2008). In heat stress studies, the *GAPDH* gene was also found to be the most suitable reference gene in pigeonpea (Sinha et al. 2015) and *Lilium regale* (Liu et al. 2016) whereas this gene was not a suitable reference gene in cicer (Reddy et al. 2016) and maize (Lin et al. 2014). The *ACT 7* gene was not the suitable reference gene in *Caragana intermedia* (Zhu et al. 2013) for qRT-PCR studies in heat stress condition. According to our results, *GAPDH* and *ACT 7* genes were the most suitable reference genes for normalizing the gene expression data in guar under heat stress condition. Our findings are expected to help in the identification of genes involved in heat tolerance in guar.

Guar is generally cultivated in soils which have elevated salt levels. Though guar is moderately tolerant to salinity but its productivity is reduced under high salt stress conditions. Our studies showed that *GAPDH* and *EF-1a* genes were the suitable reference genes for gene expression studies in guar under salt stress condition. In the previous studies on salinity stress, the *GAPDH* gene was found to be most suitable reference gene in pigeonpea (Sinha et al. 2015) and *Platycladus orientalis* (Chang et al. 2012) and the *EF-1a* gene was most appropriate in maize (Lin et al. 2014), *Caragana intermedia* (Zhu et al. 2013) and potato (Nicot et al. 2005). In contrast, the *GAPDH* gene was found to be least stable in rice (Jain et al. 2006) and *Brassica juncea* (Chandna et al. 2012) whereas the *EF-1a* gene was not a suitable reference gene in *Platycladus orientalis* (Chang et al. 2012) under salt stress condition. On the basis of our results, now it will be possible to design experiments to identify guar genes involved in tolerance to salt stress.

In Northern India and Pakistan, guar is mainly cultivated once in a year by sowing in the month of July. It can also be cultivated for second time in a year by sowing at the end of winter in the month of February/March. For this purpose, guar varieties that can tolerate cold during initial stages of growth will be required. Our study showed that *TUA* and *UBC 2* genes were most suitable as a reference for accurate normalization of the gene expression data under cold stress condition in guar. In the previous reports, *TUA* and *UBC 2* genes were found to be most suitable reference genes under cold stress study in *Salicornia europaea* (Xiao et al. 2015), whereas the *UBC* 2 gene was the most suitable for cold stress study in *Platycladus orientalis* (Chang et al. 2012). The *TUA* and *UBC 2* genes were not the proper reference genes in cold stress condition in potato (Nicot et al. 2005), maize (Lin et al. 2014), *Caragana intermedia* (Zhu et al. 2013) and radish (Duan et al. 2017). Our results will be helpful to the studies of gene expression analysis to identify cold tolerance genes of guar.

The *TUB* gene in our study was the unsuitable reference gene in tissue set and cold stress set whereas *TUA* and *EF-1α* genes were the most unsuitable reference genes in seed development and nitrogen stress sets, respectively. The *18S rRNA* was the most unsuitable reference gene in drought, heat and salt stress sets.

The validation of four selected reference genes was done by normalizing the expression of *CtMT1* gene which plays an important role in plant development and stress response. The results obtained with the most stable reference gene were found to be in agreement to the results of the expression of *CtMT1* gene in the tissues of other plants, however, the results using the least stable reference genes were contradictory to the findings of previous workers (Hudspeth et al.1996; Yang et al. 2009; Ahn et al. 2012; Yang et al. 2015).

The above discussion shows that our results are in line with those of the previous studies that no single gene is a suitable reference gene under different conditions in different plant species. Therefore, it is imperative to select a suitable reference gene for normalizing the qRT-PCR gene expression data in a particular species for each experimental condition. Our findings of suitable reference genes for various conditions in guar are expected to provide a boost to the gene expression studies in this crop. Such studies will considerably improve our understanding of the molecular mechanisms of stress tolerance and gum synthesis in guar and will be helpful in developing improved varieties of this industrially important crop.

## Author contributions

PSJ and NK designed the experiments, did all the experimental work, analysed the data and wrote the manuscript. GSR designed the experiments, interpreted the data and wrote the manuscript. All authors read and approved the final manuscript.

## Acknowledgments

We would like to thank Prof. S. S. Dudeja for his help in arranging the seeds of the guar variety HG-365 and Dr. Harsh Chauhan, Department of Biotechnology, IIT Roorkee for his valuable suggestions regarding gene expression studies. Financial support in the form of assistantships to PSJ and NK by the Ministry of Human Resource Development, Govt. of India, is gratefully acknowledged.

## Funding

This research was not supported by any research grant.

## Footnotes

Availability of data and supplemental materials

The supporting data for this article have been added as supplemental materials and uploaded to figshare. The partial cDNA sequences of nine house-keeping genes of *C. tetragonoloba* obtained during this work have been submitted to NCBI but these sequences have not been made public yet. The NCBI accession numbers of the submitted partial cDNA sequences are MF370605 *(ACT 11)*, MF370606 *(ACT 7)*, MF370607 *(TUA)*, MF370608 *(TUB)*, MF370609 *(GAPDH)*, MF370610 *(UBQ 10)*, MF370611 *(EF-1α)*, MF370612 *(UBC 2)* and MF370613 *(CYP)*.

## Supplementary Figure S1 Agarose gel electrophoresis of PCR products of candidate reference genes

The loading of the PCR products in the agarose gel was as follows: 1-*18S rRNA* (113 bp), 2-*EF-1α* (180 bp), 3- *TUA* (205 bp), 4 – *TUB* (167 bp), 5- *UBC 2* (148 bp), 6- *ACT 7* (161 bp), 7- *ACT 11* (154 bp), 8- *UBQ 10* (197 bp), 9- *GAPDH* (192 bp) and 10- *CYP* (93 bp). M- 100 bp DNA ladder.

